# Investigating the potential for mirror self-recognition in honeybees (*Apis mellifera*): a preliminary study using the mark test

**DOI:** 10.1101/2025.10.14.682345

**Authors:** Gentaro Shishimi, Rena Akaishi, Mana Matsukami

**Author notes:** (GS). These authors contributed equally to this work (Rena Akaishi, Mana Matsukami).

## Abstract

Mirror self-recognition (MSR) is a key indicator of self-awareness, primarily studied in vertebrates. Investigating MSR in invertebrates such as the western honeybee (*Apis mellifera*), an insect with advanced cognition, is essential to understanding the evolutionary origins and potential convergent evolution of self-recognition. We conducted two experiments using a classic mark test, applying a paint mark to the clypeus, a body part invisible to the bee without a mirror, and compared head-directed grooming between bees with and without mirror access. In Experiment 1 (n=12), without prior mirror exposure, bees did not increase their total grooming activity, although they did groom for a significantly higher proportion of time while facing the mirror. In Experiment 2 (n=9), bees with prior mirror exposure showed significantly higher total grooming activity than controls, an effect driven by an increase in grooming while facing the mirror. However, the critical test for MSR, a direct comparison of grooming while facing the mirror versus facing away from it, was not significant in either experiment’s mirror group. Our findings suggest the observed behaviors are not indicative of MSR. Instead, they are better explained by simpler mechanisms, such as an orienting response to a novel stimulus (Exp 1), generalized agitation from a perceived social stimulus (Exp 2), or tactile irritation from the mark itself, an interpretation consistent with findings in other invertebrates. This exploratory research, limited by small sample sizes, provides preliminary yet inconclusive evidence. We discuss our results in the context of key methodological prerequisites for the mark test, such as mark salience, and the broader possibility that MSR-like behaviors reflect a learned perceptual-motor skill rather than an abstract self-concept. Thus, this study contributes not by providing a definitive answer, but by highlighting these critical challenges and establishing a rigorous framework for future investigations designed to disentangle these alternative hypotheses.

## Introduction

The capacity for self-recognition, the ability to identify oneself as a distinct entity separate from the environment and others, has long been a central topic in cognitive science. Mirror self-recognition (MSR), typically assessed using the mark test, is a primary experimental benchmark for this capacity in non-human animals and is widely regarded as a key indicator of self-awareness and potentially other higher-order cognitive functions [1, 2]. Since Gallup’s seminal study in 1970, which first provided experimental evidence of MSR in chimpanzees [3], this phenomenon has attracted substantial interest across multiple disciplines, including primatology, comparative psychology, and neuroscience. This classic mark test involves applying a mark to a body part invisible to the subject without a mirror and observing whether self-directed behaviors (e.g., touching, inspecting) of the marked area increase in the presence of a mirror compared to its absence [1, 2].

Although MSR was initially considered a hallmark of great apes and humans, a large-scale study on chimpanzees revealed that MSR is a developmental phenomenon, not universally present in all individuals [4]. Subsequent research has reported MSR or MSR-like behaviors in a select group of other mammals and birds, including bottlenose dolphins [5], Asian elephants [6], and Eurasian magpies [7]. These species often, though not exclusively, possess relatively large brain capacities and exhibit complex social structures [8, 9]. However, the interpretation of these findings, particularly in non-primate species, remains a subject of considerable debate and methodological scrutiny [10, 11]. Recent research has further explored the phylogenetic breadth of MSR, questioning’how simple a nervous system can support MSR?’. In this context, the success reported in the cleaner wrasse (*Labroides dimidiatus*), a fish species [12], received significant attention by suggesting that MSR capability is not necessarily limited to large brains or specific lineages, although this interpretation itself remains highly contested and subject to alternative explanations [13, 14]. Furthermore, studies in mice have shown that the activation of hippocampal neuronal ensembles by visuotactile integration promotes mirror-induced self-directed behavior, suggesting a role for the hippocampus in MSR in mammals [15].

Evidence for MSR in invertebrates is particularly limited and often debated. For instance, a study on three ant species (*Myrmica sabuleti*, *Myrmica rubra*, and *Myrmica ruginodis*) reported mark-directed behavior in adults but not in very young subjects, requiring cautious interpretation [16]. Research on paper wasps (*Polistes metricus*) found increased exploratory behaviors towards mirrors but did not yield conclusive evidence of MSR [17].

Among invertebrates, arguably the most extensively studied group for complex cognition is the cephalopods (octopus, squid, and cuttlefish). However, a recent comprehensive review concluded that evidence for mirror self-recognition (MSR) in cephalopods remains weak and contested [18]. While some species like the oval squid show strong exploratory behaviors such as touching their mirror image [19], a preliminary attempt to apply the mark test to the common octopus found that the observed mark-directed grooming was likely triggered by the tactile sensation of the mark itself, rather than visual self-recognition [20].

Currently, the ghost crab (*Ocypode quadrata*) is reported as potentially one of the phylogenetically lowest organisms showing evidence of MSR [21], though again, interpretation requires careful consideration of alternative explanations.

This approach, focusing on an invertebrate, directly addresses a foundational critique within comparative psychology. For decades, influential voices have called for the investigation of fundamental cognitive processes at widely separated points in the phylogenetic scale to truly test the generality of psychological principles, arguing against the field’s historical over-reliance on a few mammalian models [22]. Against this complex and evolving background, the present study investigates MSR in the western honeybee (*Apis mellifera*). The honeybee is an insect phylogenetically situated between crustaceans and vertebrates and is renowned for its remarkable cognitive abilities despite possessing a brain containing only about one million neurons [23]. Honeybees exhibit a surprisingly diverse suite of complex cognitive functions. Indeed, a long tradition of research has systematically analyzed honeybee learning by drawing direct comparisons to vertebrate performance, establishing the honeybee as a powerful model for the comparative analysis of cognition [24, 25]. This comparative work has been extensively reviewed in the context of visual cognition and perception [26, 27].

Their capacity for concept learning is particularly notable [28]. For instance, their ability to learn the abstract concepts of’sameness’ and’difference’ has been reported [29, 30], and the topic has been recently reviewed [31]. A key factor underpinning this ability is a robust visual working memory. While an early study reported this memory to last for 6.5 seconds [32], subsequent research has demonstrated that it can, under certain conditions, persist for approximately 2.5 minutes [30]. They can also learn to apply two abstract concepts simultaneously [33], learn concepts of oddity/nonoddity [34, 35], relative size [36], and above/below spatial relationships [37]. Their cognitive suite further includes maze navigation [38], numerical cognition including understanding zero and parity [39, 40], perception of visual illusions [41], and flexible behavioral strategies in learning tasks [42]. This suite of abilities has also inspired research in robotics and artificial intelligence, such as the development of neuromorphic vision systems [43]. They also exhibit emotion-like states interpreted as “pessimistic decision-making” [44]. Furthermore, they demonstrate sophisticated decision-making strategies, such as selectively “opting out” of difficult discrimination tasks, a behavior that has been rigorously debated in the context of uncertainty monitoring or metacognition [45].

In social contexts, evidence suggests sophisticated social cognitive capabilities, such as learning from conspecifics using visual cues [46], as well as nestmate recognition using olfactory and tactile cues [47], and remarkably, they have even been reported to discriminate human faces [48]. Such complex visual recognition is likely important for recognizing conspecifics [49] and adjusting behavior accordingly, particularly during foraging [50].

Particularly relevant to MSR is the honeybee’s capacity to process symmetrical visual information. Previous work by Giurfa et al. (1996) has shown that honeybees can extract bilateral symmetry as a visual feature [51], and furthermore, Stach & Giurfa (2001) demonstrated that they can generalize learned visual patterns to their mirror images [52]. This suggests a basic ability to process mirror image information, a potential prerequisite for MSR. However, their study focused on generalization in pattern recognition, a different cognitive process from recognizing one’s own image in a mirror (MSR). The present study aims to test whether this mirror image processing ability extends to the recognition of the self-image using the mark test.

Investigating MSR in honeybees holds significant implications. Firstly, if MSR is confirmed in an insect with a relatively simple nervous system, it would strongly suggest that higher cognitive functions like self-recognition do not necessarily require large brains or specific structures (e.g., neocortex), thereby deepening our understanding of the evolutionary origins and neural basis of self-recognition [8]. It might eventually open possibilities for identifying the neural circuits involved in MSR within the relatively tractable honeybee nervous system [23, 24]. Secondly, findings on honeybee MSR, highlighting complex cognition within a relatively simple nervous system [24], could contribute not only to psychology and biology but also inform robotics and artificial intelligence research [53]. For example, computational approaches have already demonstrated the feasibility of functionally imitating honeybee concept learning processes with simple neural networks inspired by the insect brain [54], as well as with other computational frameworks such as Vector Symbolic Architectures [55], potentially shedding light on the foundations required for capabilities like self-recognition.

We chose honeybees over paper wasps for several reasons. First, honeybees exhibit a higher degree of social complexity [56]. This is a key consideration, as a prominent theory for the evolution of self-recognition, the social intelligence hypothesis, posits that the cognitive demands of complex social life are a primary driver for the evolution of self-awareness [9, 57, 58]. Indeed, strong evidence from paper wasps suggests that selection for individual recognition can lead to the evolution of specialized cognitive abilities, such as dedicated face learning [59]. This proposed link between sociality and mirror-directed behavior has also been observed in cephalopods. For example, highly social species (squid) have been shown to have strong interest in their mirror image, whereas asocial species (octopus) tend to show little or no reaction, suggesting that the cognitive pressures of social life may be a key factor in the evolution of such behaviors [60]. Therefore, highly social insects like honeybees represent a critical test case for the taxonomic limits of this hypothesis. Honeybee colonies comprise tens of thousands of individuals (compared to tens or hundreds in independent-founding paper wasps) and feature distinct morphological and behavioral caste differentiation (queen, workers, drones) [47, 56]. They have developed sophisticated social systems, including advanced communication via the waggle dance [47] and collective defense behaviors like heat-balling [61]. Such a complex social environment likely imposes strong selection pressure on the ability to accurately discriminate self from others (nestmates, non-nestmates, predators, etc.) and adjust behavior accordingly, potentially favoring self-related cognitive abilities. Efficient functioning of a system where tens of thousands live closely, dividing and coordinating diverse tasks (e.g., foraging, brood care, nest construction/maintenance/defense, cleaning, etc.), arguably requires individuals to differentiate, to some extent, their own state and actions from those of others. Mistaking reflections (e.g., from water surfaces or dew) during activities outside the hive could be maladaptive, wasting energy and time [62]; thus, the ability to distinguish their own reflection from other individuals could be adaptive. Second, as mentioned earlier, various cognitive abilities potentially related to MSR expression, such as the ability to recognize and learn from conspecifics using visual cues [46], complex visual pattern recognition [48], mirror-image generalization [52], and concept learning [29–31, 33–37, 42, 43, 50], have already been reported in honeybees, suggesting a higher potential for MSR presence compared to species where these prerequisite abilities are less documented.

It is noteworthy that previous studies have used mirrors (on the floor of experimental tunnels or during flight over water) to investigate how honeybees use optic flow from the ground for altitude control [62, 63]. These studies examined flight behavior changes when visual cues were manipulated using horizontal mirrors and differ significantly in purpose (investigating visually guided flight, not MSR via self-directed behavior facing a vertical mirror) and experimental setup from the present study. This study aimed to determine whether honeybees exhibit mirror self-recognition (MSR) by applying the classic mark test. Furthermore, by exploring a species with a miniature nervous system, this research also addresses the important call to investigate the cognitive limits of animals, which is as crucial as demonstrating their capabilities for understanding the diversity and evolution of cognition [64]. We hypothesized that if honeybees possess MSR capabilities, subjects exposed to a mirror would show increased self-directed grooming towards a mark placed on their head, compared to control subjects without a mirror. To investigate this hypothesis, we compared the frequency and duration of head-directed grooming directed at a marked area on the head between bees presented with a mirror (mirror group) and those without (no-mirror/sham mirror group). By applying the same classic paradigm to a phylogenetically distant invertebrate, this study also provides an opportunity to explore the potential for the convergent evolution of complex cognitive traits [22, 25, 65] such as self-recognition. Furthermore, to examine the effect of prior mirror experience, we conducted Experiment 1 (without pre-exposure) and Experiment 2 (with pre-exposure).

## Materials and methods

### Animals

Foraging worker bees (*Apis mellifera*) were collected as needed from hives on the rooftop of Mita International School of Science (Tokyo, Japan) as they departed from the hive entrance. Collected bees were therefore foragers, though precise age was not controlled. The sample sizes were smaller than initially planned due to colony collapses during the experimental periods, which is a notable limitation of this study.

### Experiment 1: Mark test (no prior mirror exposure)

#### Subjects

Thirteen worker bees were subjected to the experimental procedure. They were randomly assigned to the Mirror group (n = 7; Ss 1, 2, 5, 6, 9, 10, 13) or the Control group (n = 6; Ss 3, 4, 7, 8, 11, 12). (As noted later and in the Results section, the final number of subjects analyzed was reduced due to marking failures, poor recovery from anesthesia, or abnormal behavior during the experiment.)

#### Apparatus

Transparent acrylic boxes served as individual test boxes for observation. For subjects up to S8, the box size was 13.5 cm L × 9.7 cm W × 7.1 cm H; for S9 onwards, a slightly smaller box (10.0 cm L × 10.2 cm W × 4.6 cm H) was used. This change was implemented to improve visibility for subsequent video analysis. A horizontal platform served as the base for the test box, and the platform was enclosed by white paper (overall enclosure size: approx. 52 cm L × 55 cm W × 38 cm H) to block external visual disturbances and avoid reflections on the test box (Fig 1).

**Fig 1.**
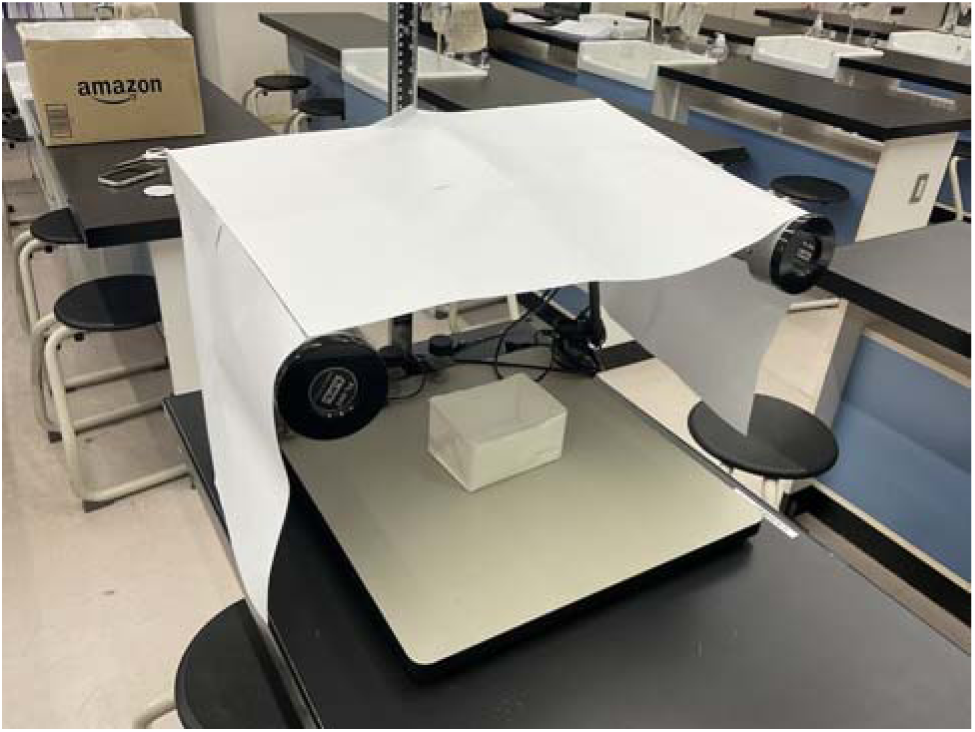
Experimental setup for Experiment 1, showing a test box on a horizontal platform, enclosed by white paper to block external visual disturbances and avoid reflections on the test box.

#### Marking procedure

Approximately 10 bees were captured near the hive entrance in a clear plastic bag. To induce a temporary hypothermic state, the bag containing the bees was placed in a freezer (approx.-18°C) until they became immobile (approx. 4 min 30 s). The anesthetized bees were then removed from the bag and each was individually marked on the clypeus. A small amount of white correction fluid (Pentel Co., Ltd. XEZL1-W) was applied to the tip of a toothpick and used to place a small mark on the bee’s head, specifically on the clypeus below the compound eyes and above the mandibles (Fig 2), adopting a similar mark placement to that employed in tests with paper wasps [17]. This marking procedure presented technical challenges due to the small size of the bees and their rapid arousal. A number of failures occurred, such as incorrect mark size or placement, or application of fluid to unintended areas (e.g., antennae, mouthparts).

**Fig 2.**
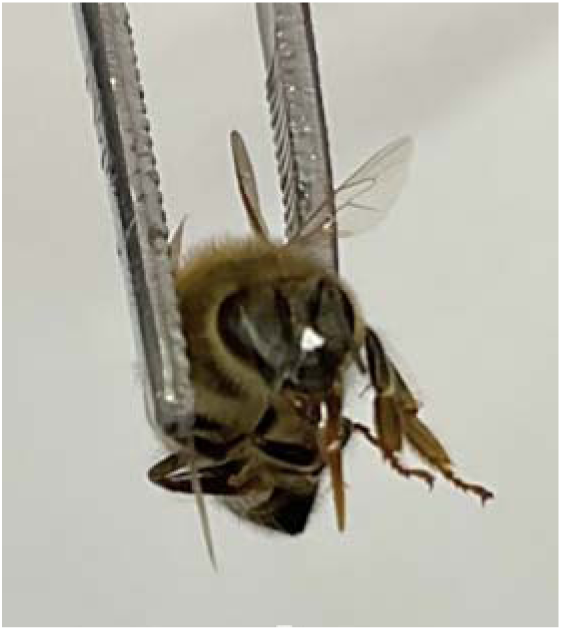
Honeybee (*Apis mellifera*) with a white mark applied to the clypeus (below the compound eyes and above the mandibles) for the mark test.

Subjects judged to have been improperly marked were excluded before proceeding. This cooling procedure, a common method for temporarily immobilizing honeybees for manipulation, was used to minimize invasiveness while allowing for accurate marking. However, it is important to acknowledge that the procedure itself, particularly the potential for physical irritation or olfactory cues from the marking fluid, could influence subsequent behavior and thus requires careful interpretation of the results.

#### Experimental procedure

After the marking procedure, bees were allowed to recover. Once a subject began moving, it was fed a small amount of 50% (w/w) sucrose solution via pipette to provide energy. Each bee was then placed individually into a transparent test box enclosed by white paper on all sides except the top. For each trial, the test box containing the individual that appeared most physically intact and exhibited the most normal recovery behavior was selected. This selected box was placed onto a horizontal platform, and a 5-minute acclimatization period commenced when the bee inside started walking spontaneously. After acclimatization, one of two conditions was applied. For the Mirror group, an acrylic mirror (commercially available, cut to fit the side) was inserted vertically along one of the side walls of the test box; specifically, a short side wall for the larger rectangular box (subjects up to S8) and the slightly longer side wall for the smaller box (subjects from S9 onwards). For the Control group, subjects up to S8 received no mirror. For subjects S9 onwards, a’sham mirror’ (the back of a mirror covered with non-reflective white cardstock) was inserted to control for the potential effects of the insertion procedure and to provide a non-reflective visual stimulus, ensuring similar physical manipulation between groups. Behavior was video-recorded from directly above the test box using an iPad (Apple Inc.) for 30 minutes following the acclimatization period. Note that during recovery from anesthesia, a few subjects died or exhibited prolonged immobility, and these were excluded from the experiment.

### Experiment 2: Mark test (prior mirror exposure)

#### Purpose

Given that Experiment 1 yielded no clear MSR evidence, and considering the importance of prior mirror experience often highlighted in MSR research [2], Experiment 2 aimed to test the effect of pre-exposure under more controlled conditions (consistent sham mirror control, extended observation time, uniform box size).

#### Subjects

Nine new worker bees were subjected to the procedure. They were randomly assigned to the Mirror group (n = 5; Ss 1, 2, 4, 6, 9) or the Sham Mirror group (n = 4; Ss 3, 5, 7, 8). (As in Experiment 1, the number analyzed reflects exclusions due to the criteria mentioned above).

#### Apparatus

To maintain consistency and improve visibility during video analysis, the same smaller transparent acrylic boxes (10.0 cm L × 10.2 cm W × 4.6 cm H) were used for initial capture, the pre-exposure session, and as test boxes for observation. The indoor setup for Experiment 2 consisted of a cardboard box (32 cm L × 40 cm W × 26 cm H) with an open top and lined entirely with white paper on the interior walls to reduce reflections and provide a uniform background. The cardboard box was placed on a horizontal platform, and a portion of the box top was covered with white paper to further minimize reflections. Additionally, one or two white paper enclosures, each used in Experiment 1 to cover a test box, were placed inside the cardboard box.

#### Pre-exposure

Pre-exposure was conducted in two stages: at the hive and subsequently in the laboratory.

1. **At the hive.** A few days before starting to collect experimental subjects, a large mirror (reflective surface: 28.5 cm × 38.5 cm) was placed in front of the entrance of the target colony’s hive (Fig 3). This provided bees entering and leaving the hive with natural exposure opportunities to their moving reflections. This procedure was intended to allow for potential associative learning between the bees’ own movements and the movements observed in the mirror reflection, which might facilitate recognition of the mirror image as their own in a more natural context, but its effectiveness remains speculative. Additionally, the exact duration and degree of attention paid by each individual were not recorded or controlled.
2. **In the laboratory.** The second stage of pre-exposure was conducted in the laboratory to habituate the bees to the presence of a mirror within the testing context, thus minimizing responses to the mirror as a novel object during the subsequent experimental phase. Bees were individually captured in the experimental boxes in front of the hive with the large mirror. To eliminate the potential impact of the mirror insertion process on the bees’ behavior, these experimental boxes, each containing a bee, were then directly placed inside the white paper enclosures with a mirror pre-installed on one side, within the indoor experimental setup for an additional pre-exposure session in the lab. Bees in both the Mirror and Sham Mirror groups had a mirror placed along one side wall (within the experimental box). We aimed for counterbalancing; however, the small sample sizes limited its full implementation. The mirror position remained consistent for each individual between the pre-exposure and testing periods. They were allowed to explore freely for 20 minutes.

**Fig 3.**
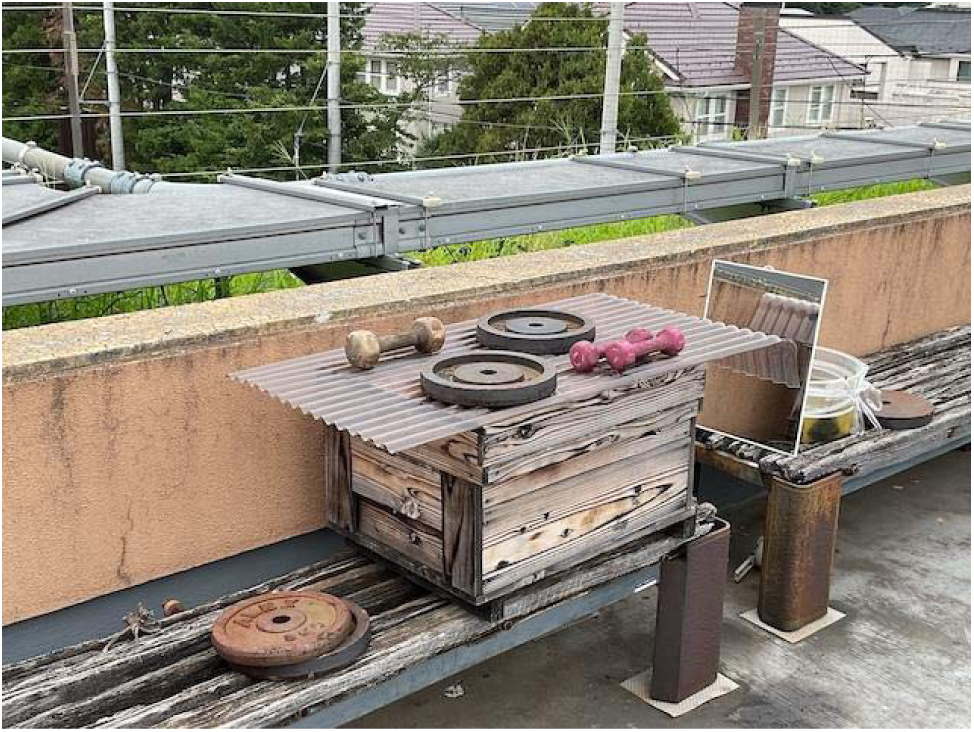
A large mirror (reflective surface: 28.5 cm × 38.5 cm) for Experiment 2 pre-exposure, placed in front of the colony hive.

#### Experimental procedure

The overall procedure for Experiment 2 generally followed that of Experiment 1, but the observation time was extended to 40 minutes to gather more behavioral data. The following steps were performed after the pre-exposure phase:

1. After the 20-minute pre-exposure session, bees were removed and anesthetized by cooling as in Experiment 1 (approx. 4 min 30 s).
2. Marking was performed as in Experiment 1. Exclusions due to marking failures occurred here as well.
3. Upon recovery, bees were individually fed 50% (w/w) sucrose solution and placed in transparent, clean test boxes. To avoid any behavioral changes potentially induced by inserting either the mirror or the sham mirror during the experiment, the experimental setup already included individual white paper enclosures, each with either a mirror (Mirror group) or a sham mirror (Sham Mirror group) pre-installed along one longer side. Then, one or two transparent test boxes, each containing a bee, were placed inside the corresponding white paper enclosures for each trial.
4. The bee in its test box was observed, and its behavior was video-recorded for 40 minutes starting from when it began walking spontaneously. In Experiment 2, in order to observe the effects of prior mirror experience in more detail, unlike Experiment 1, video recording was started immediately after the bee began walking spontaneously. The aim was to capture a wider range of behavioral changes by including the behavior immediately after recovery from anesthesia. To improve efficiency, two subjects were sometimes tested concurrently in separate boxes and their respective white paper enclosures, but they remained visually and physically isolated from each other. Subjects showing poor recovery from anesthesia were excluded from analysis.

### Data collection and behavioral analysis

After the experiment, recorded videos were observed by two independent observers (RA and MM, the second and third authors, who were also the experimenters). To ensure consistency and assess inter-observer reliability, all video analysis was conducted by the two observers (RA and MM). This process included obtaining measurements from videos played back at slow speed as needed to reliably capture fast movements. The following behavioral metrics were measured.

#### Grooming

Grooming was defined as any action in which the forelegs touched or scrubbed any part of the head region, including the clypeus, compound eyes, and antennae. During analysis, it was not possible for observers to reliably distinguish between actions specifically directed at the paint mark and more general grooming actions directed at nearby areas, such as the antennae. Therefore, all recorded grooming data represent general head-directed grooming rather than exclusively mark-directed actions. Total (T) frequency and duration (in seconds, measured with 0.01 second precision) were recorded. Due to the fast movements of some grooming behaviors, which made accurate measurement difficult at normal playback speed, videos were slowed down as needed for reliable measurement.

#### Body Orientation during Grooming

The bee’s body orientation relative to the mirror (or sham mirror/hypothetical mirror location for some controls) was recorded at the moment grooming occurred.

Orientations were coded as Facing (F) if the bee’s body axis was within approximately ±90° of perpendicular to the mirror surface; Away (A), with the body axis outside the ±90° range; or Not facing or away (N), for orientations that were roughly sideways or ambiguous (Fig 4). For Control group subjects up to S8 in Exp 1, a mirror position was randomly assumed to determine orientation. For all subsequent control and sham mirror subjects, the inserted sham mirror served as the reference.

**Fig 4.**
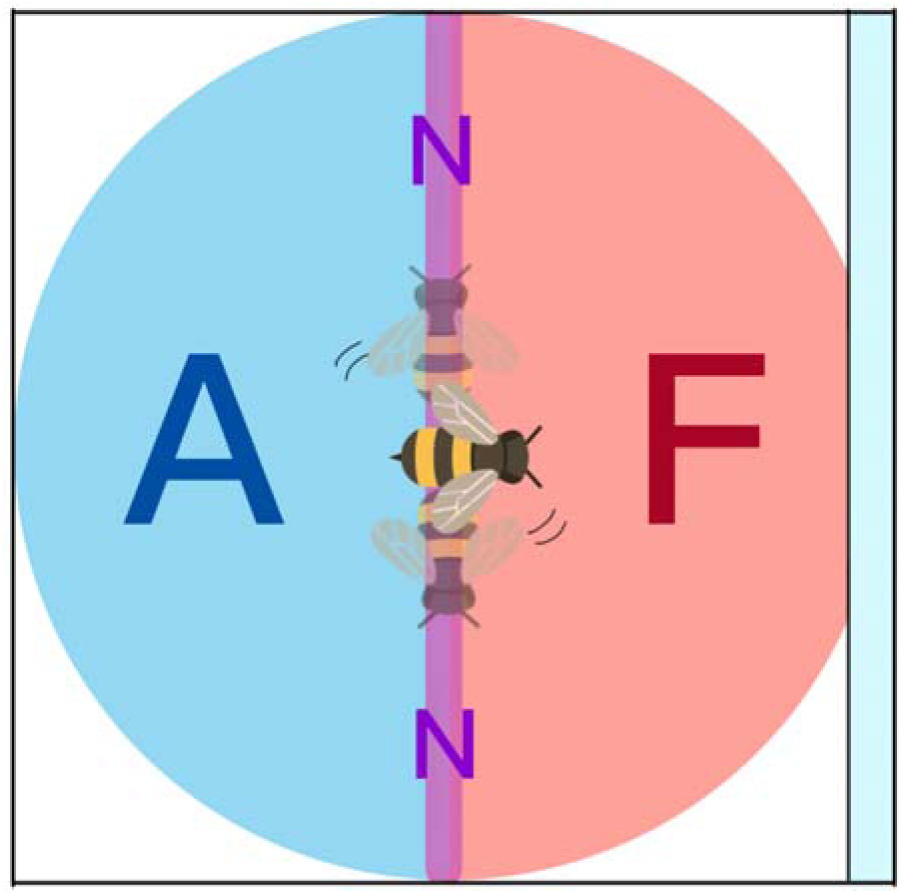
Classification of honeybee body orientation relative to the mirror during grooming.* * F: Facing the mirror, A: Away from the mirror, N: Not facing or away.

The “N” category was included to avoid forcing ambiguous observations into the “F” or “A” classifications, thereby ensuring a more rigorous application of these criteria.

However, data for the “N” category proved to have low inter-observer reliability (Pearson’s r:-0.232 to 0.968), likely due to its infrequent occurrence. Therefore, these data were excluded from the main behavioral analyses and are presented only as supplementary information. The main statistical analyses focused on the more reliable Total (T), Facing (F), Away (A), and Facing/Total ratio (F/T) metrics.

## Statistical analysis

For each individual, the values measured by the two observers were averaged for analysis. Inter-observer reliability was assessed using Pearson’s correlation coefficient (Pearson’s r) for each metric (T, F, A, N frequency and duration). Consistency between duration and frequency metrics was also assessed using Pearson’s r. Between-group comparisons (Mirror vs. Control/Sham Mirror) for T, F, A, and N grooming frequency and duration were performed using Welch’s t-test, selected to accommodate potentially unequal variances and small sample sizes. The proportion of mirror-facing grooming (F/T, where T = F+A+N) was calculated as an index of directional bias and compared between groups (Welch’s t-test). Within-group comparisons (F vs. A frequency and duration) in the Mirror group were performed using paired t-tests. The statistical significance level was set at α =.05. Given the small sample sizes, exact p-values are reported, and effect sizes (Cohen’s d) are provided for context, but all statistical results should be interpreted with caution due to limited power.

### Individual analysis (binomial test)

To examine individual variation in addition to group-level trends, data from each observer for each individual were analyzed separately. The number of grooming bouts Facing (F) versus Away (A) from the mirror/sham was compared using a two-tailed binomial test (α =.05) to assess significant directional bias. Raw count data from each observer were used as the test requires integer values.

### Ethical considerations

All experimental procedures were conducted with the highest regard for animal welfare. We aimed to minimize the number of bees used, acknowledging challenges in handling foraging bees from outdoor hives, including recovery from cooling anesthesia, marking precision, or behavior. In Experiment 1, approximately 10 bees were captured per session in a plastic bag, an approach that sometimes led to necessary exclusions. In Experiment 2, bees were introduced individually into experimental boxes, minimizing unused individuals. Cooling anesthesia, when applied, was limited to the minimum necessary (approx. 4 min 30 s) for safe handling and rapid recovery. Marking was performed swiftly to minimize stress. Bees showing prolonged immobility or abnormal behavior were immediately excluded.

## Results

### Inter-observer reliability and correlation between duration and frequency

In both experiments, high inter-observer reliability was found for all grooming behavior measurements, including total and orientation-specific frequency and duration (T, F, A) and the facing/total (F/T) ratio. In Experiment 1, Pearson’s r was ≥.960 for duration metrics and ≥.913 for frequency metrics. Similarly, in Experiment 2, r was ≥.898 for duration and ≥.918 for frequency. Strong positive correlations were also observed between duration and frequency for each metric (T, F, A, and F/T) within each experiment (Experiment 1: r ≥.984; Experiment 2: r ≥.890). This indicates that duration and frequency were closely related measures of grooming behavior, reflecting similar underlying tendencies. Therefore, we report statistical results for both metrics, as the trends were consistent across both.

### Experiment 1: Mark test (no prior mirror exposure, 30-min observation)

To visually represent the results of Experiment 1, the following box plots are provided: Fig 5 shows the grooming duration for T, F, A, and N; Fig 6 shows the F/T ratio for grooming duration; Fig 7 shows the grooming frequency for T, F, A, and N; and Fig 8 shows the F/T ratio for grooming frequency. Of the 13 bees subjected to the procedure, S13 was excluded from analysis due to abnormal behavior (e.g., prolonged immobility) during the video recording. Data from the remaining 12 bees (Mirror group n = 6, Control group n = 6) were analyzed. Between-group comparisons were performed using Welch’s t-test.

**Fig 5.**
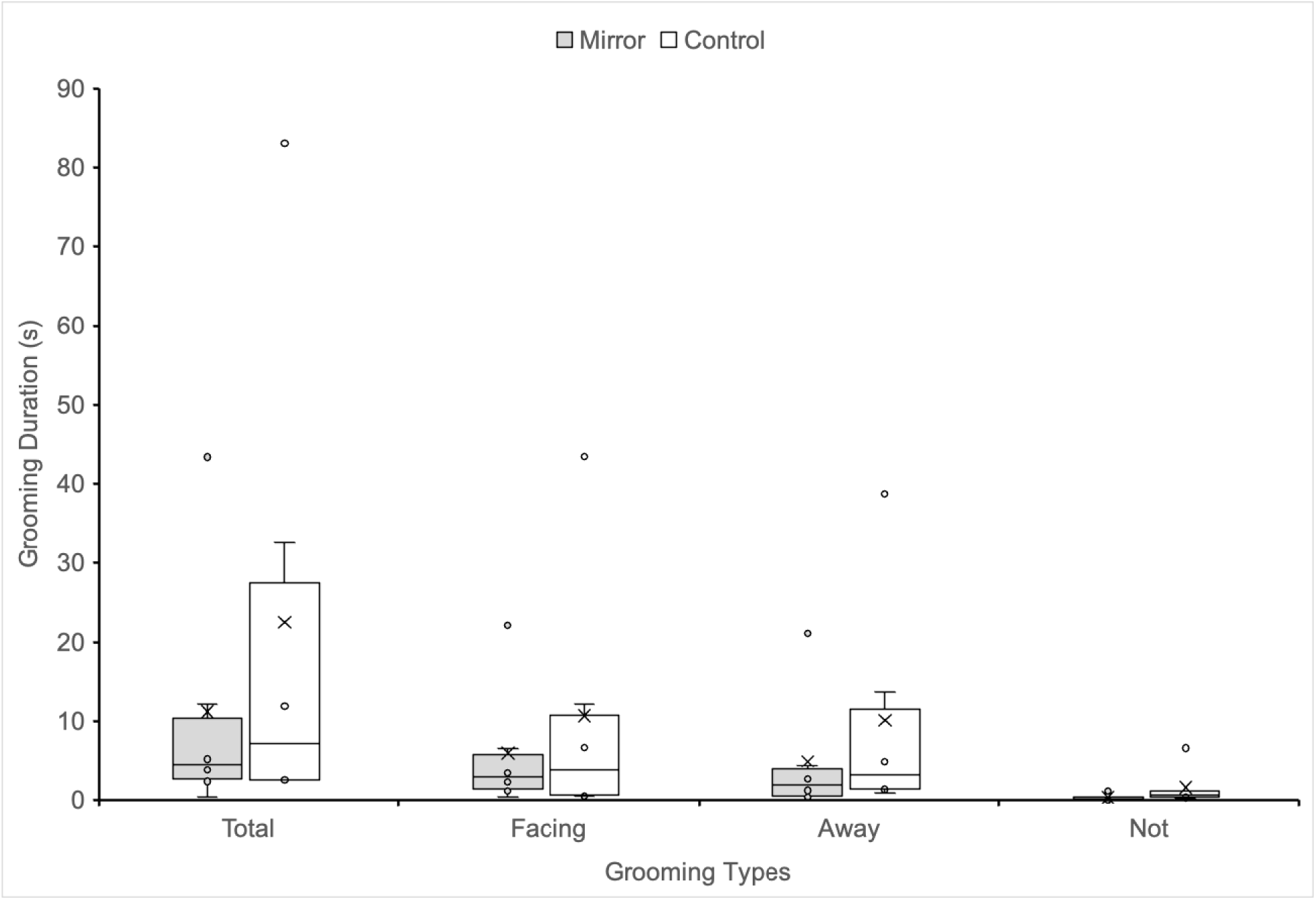
Box plots showing grooming duration (seconds) for T, F, A, and N for the Mirror group and Control group in Experiment 1. * * T: Total, F: Facing the mirror, A: Away from the mirror, N: Not facing or away.

**Fig 6.**
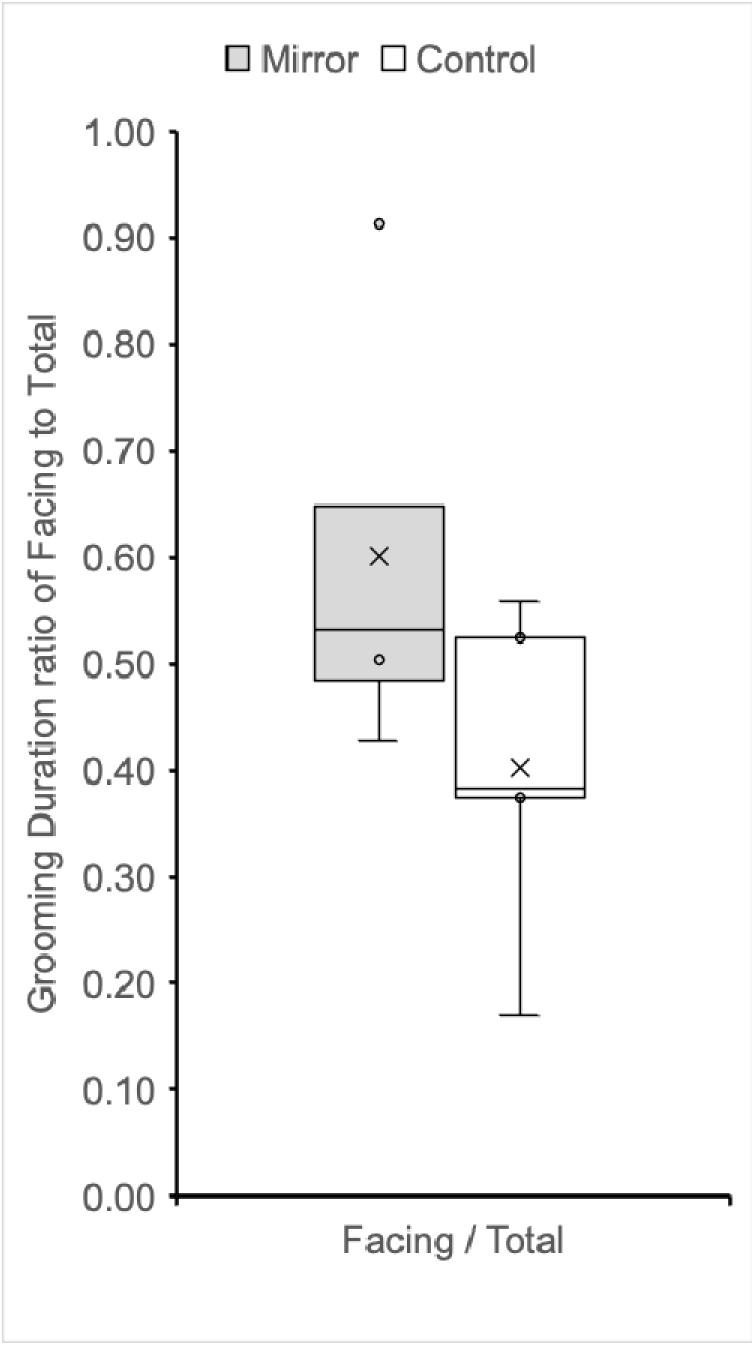
Box plots showing the F/T ratio of grooming duration for the Mirror group and Control group in Experiment 1. * * F: Facing the mirror, T: Total.

**Fig 7.**
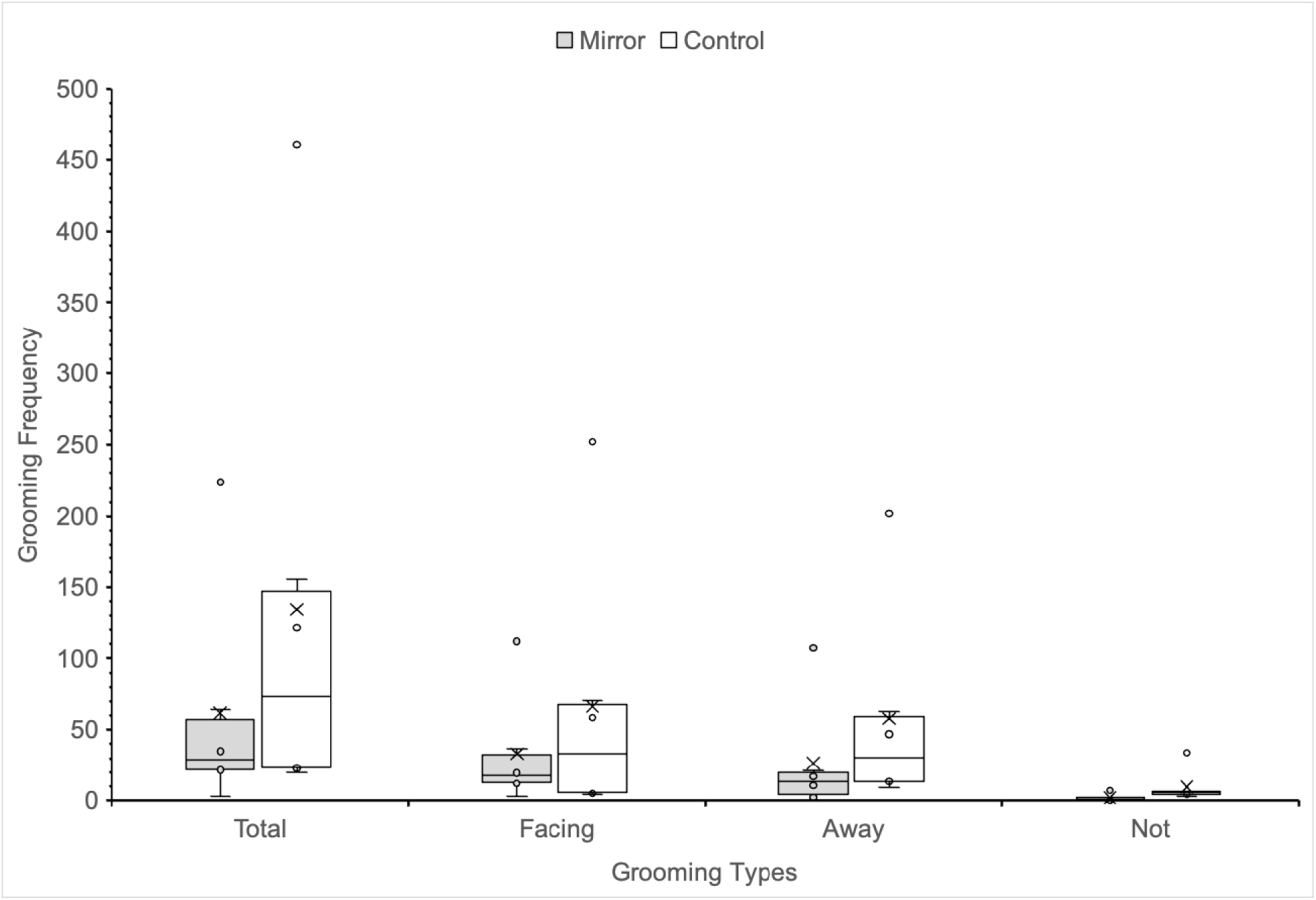
Box plots showing grooming frequency (counts) for T, F, A, and N for the Mirror group and Control group in Experiment 1. * * T: Total, F: Facing the mirror, A: Away from the mirror, N: Not facing or away.

**Fig 8.**
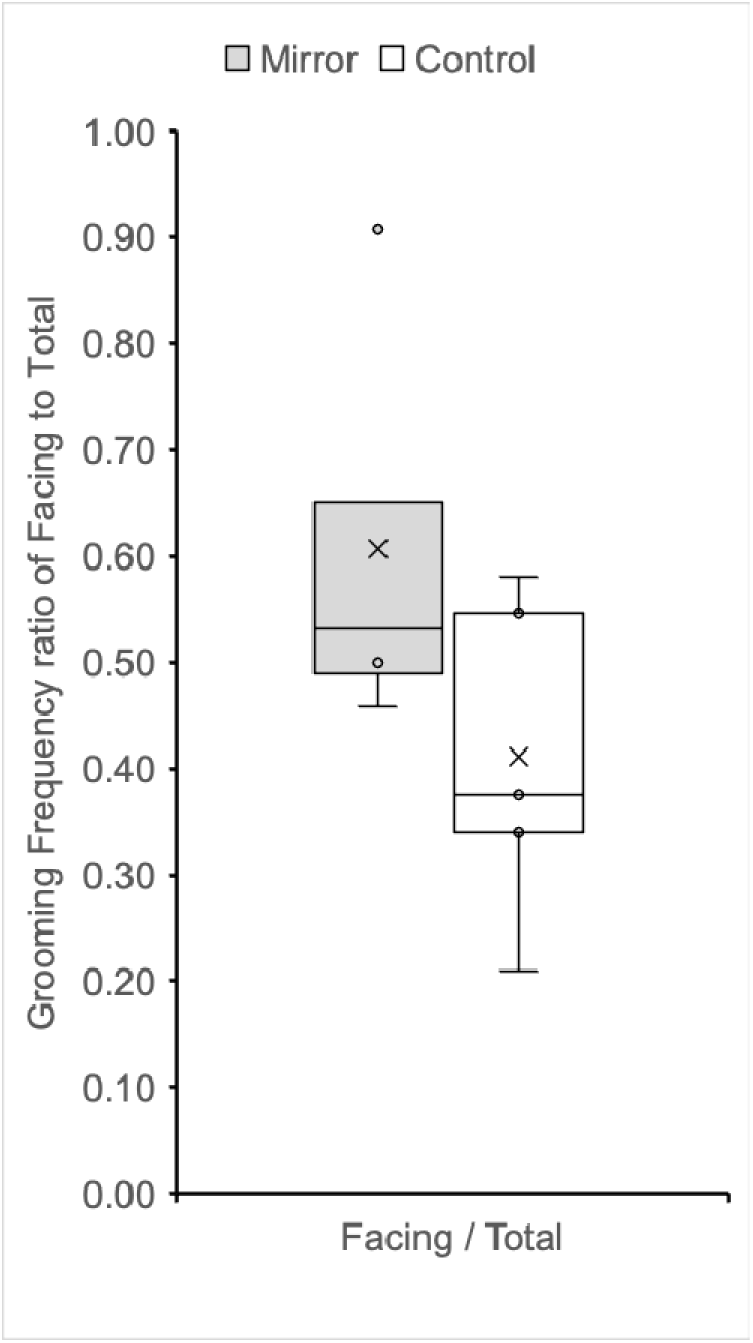
Box plots showing the F/T ratio of grooming frequency for the Mirror group and Control group in Experiment 1. * * F: Facing the mirror, T: Total.

### Grooming duration

Within-group (Mirror, n = 6): No significant difference between mean duration Facing (M = 5.997, SD = 8.219) and Away (M = 4.941, SD = 8.069) from the mirror (Paired t(5) = 1.864, p =.121, d = 0.761, 95% CI for mean difference [-0.400, 2.512], mean difference = 1.056 seconds). Within-group (Control, n = 6): No significant difference between mean duration Facing (M = 10.724, SD = 16.722) and Away (M = 10.189, SD = 14.815) from the hypothetical/sham mirror (Paired t(5) = 0.558, p =.601, d = 0.228, 95% CI for mean difference [-1.930, 3.000], mean difference = 0.535 seconds). Between-group (Total): No significant difference between Mirror (M = 11.218, SD = 16.276) and Control (M = 22.518, SD = 31.848) groups (Welch’s t(7.445) =-0.774, p =.463, d =-0.447, 95% CI for mean difference [-45.413, 22.813], mean difference =-11.300 seconds). Between-group (Facing): No significant difference between Mirror (M = 5.997, SD = 8.219) and Control (M = 10.724, SD = 16.722) groups (Welch’s t(7.283) =-0.621, p =.553, d = - 0.359, 95% CI for mean difference [-22.574, 13.119], mean difference =-4.728 seconds). Between-group (Away): No significant difference between Mirror (M = 4.941, SD = 8.069) and Control (M = 10.189, SD = 14.815) groups (Welch’s t(7.727) =-0.762, p =.469, d =-0.440, 95% CI for mean difference [-21.228, 10.732], mean difference = - 5.248 seconds). Between-group (Facing/Total Ratio): The Mirror group (M =.649, SD =.243) showed a significantly higher proportion of grooming time facing the mirror compared to the Control group (M =.371, SD =.158) (Welch’s t(8.576) = 2.354, p =.044, d = 1.359, 95% CI for mean difference [.009,.549], mean difference =.279).

### Grooming frequency

Within-group (Mirror, n = 6): A non-significant trend for more frequent grooming Facing (M = 33.167, SD = 40.163) than Away (M = 26.333, SD = 40.595) from the mirror (Paired t(5) = 2.165, p =.083, d = 0.884, 95% CI for mean difference [-1.280, 14.946], mean difference = 6.833). Within-group (Control, n = 6): No significant difference between frequency Facing (M = 66.417, SD = 95.439) and Away (M = 58.083, SD = 73.755) from the hypothetical/sham mirror (Paired t(5) = 0.865, p =.427, d = 0.353, 95% CI for mean difference [-16.427, 33.093], mean difference = 8.333). Between-group (Total): No significant difference between Mirror (M = 61.750, SD = 82.049) and Control (M =134.167, SD = 170.019) groups (Welch’s t(7.209) =-0.940, p =.378, d =-0.542, 95% CI for mean difference [-253.592, 108.759], mean difference =-72.417). Between-group (Facing): No significant difference between Mirror (M = 33.167, SD = 40.163) and Control (M = 66.417, SD = 95.439) groups (Welch’s t(6.717) =-0.787, p =.458, d = - 0.454, 95% CI for mean difference [-134.068, 67.568], mean difference =-33.250). Between-group (Away): No significant difference between Mirror (M = 26.333, SD = 40.595) and Control (M = 58.083, SD = 73.755) groups (Welch’s t(7.775) =-0.924, p =.383, d =-0.533, 95% CI for mean difference [-111.409, 47.909], mean difference = - 31.750). Between-group (Facing/Total Ratio): The proportion of grooming frequency Facing the mirror was significantly higher in the Mirror group (M =.660, SD =.232) than the Control group (M =.378, SD =.158) (Welch’s t(8.837) = 2.462, p =.036, d = 1.422, 95% CI for mean difference [.022,.542], mean difference =.282). Note on procedural variations in Exp 1 Control: Additional analyses (Welch’s t-tests) confirmed that changes in experimental box size (used for Ss up to S8 vs. S9 onwards) and the presence/absence of a sham mirror in the Control group (no mirror up to S8 vs. sham mirror S9 onwards) did not significantly affect the measured grooming metrics (ps >.05), suggesting these procedural variations did not introduce significant confounds within the limits of this small sample analysis. However, the definition of F and A for each bee in the Control group up to S8 was based on a hypothetical mirror location, which was randomly determined for each individual by a separate coin toss, whereas from S9 onwards it was based on the sham mirror.

### Summary for Experiment 1

Overall, Exp 1 showed no significant difference in the absolute amount of grooming between groups, but the Mirror group showed a significantly higher proportion of grooming frequency facing the mirror. However, since both the total grooming duration and frequency did not increase and there was no significant difference between facing and away grooming within the mirror group, the observed proportional shift provides limited support for MSR.

### Experiment 2: Mark test (prior mirror exposure, 40-min observation)

To visually represent the results of Experiment 2, the following box plots are provided: Fig 9 shows the grooming duration for T, F, A, and N; Fig 10 shows the F/T ratio for grooming duration; Fig 11 shows the grooming frequency for T, F, A, and N; and Fig 12 shows the F/T ratio for grooming frequency. Of the 9 bees subjected to the procedure, data from all 9 bees (Mirror group n = 5, Sham Mirror group n = 4) were analyzed after exclusions prior to the test phase due to marking failure, poor recovery, etc. Note that the very small sample sizes (n = 4-5 per group) significantly limit statistical power and necessitate caution in interpreting these results. Between-group comparisons were performed using Welch’s t-test.

**Fig 9.**
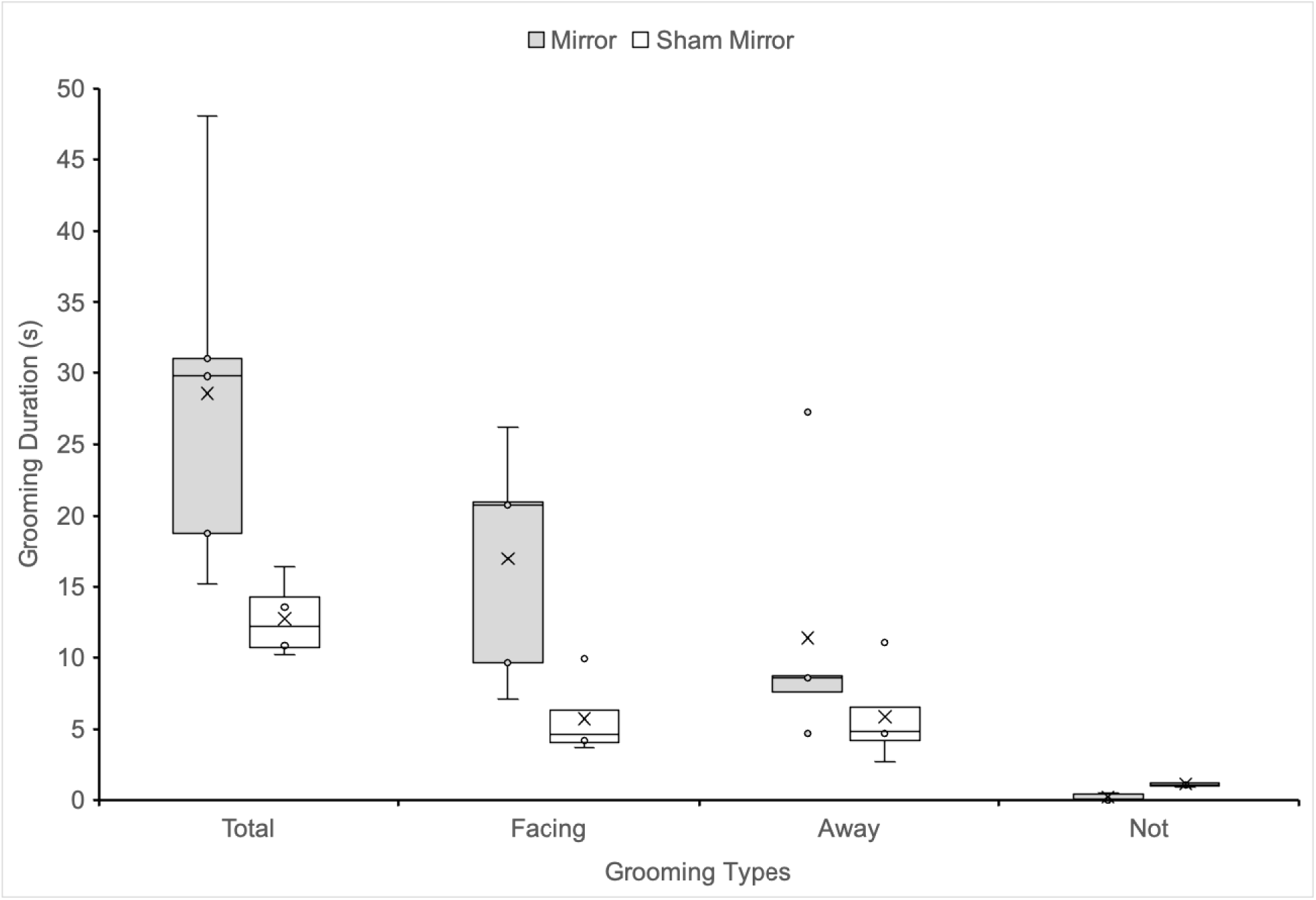
Box plots showing grooming duration (seconds) for T, F, A, and N for the Mirror group and Sham Mirror group in Experiment 2. * * T: Total, F: Facing the mirror, A: Away from the mirror, N: Not facing or away.

**Fig 10.**
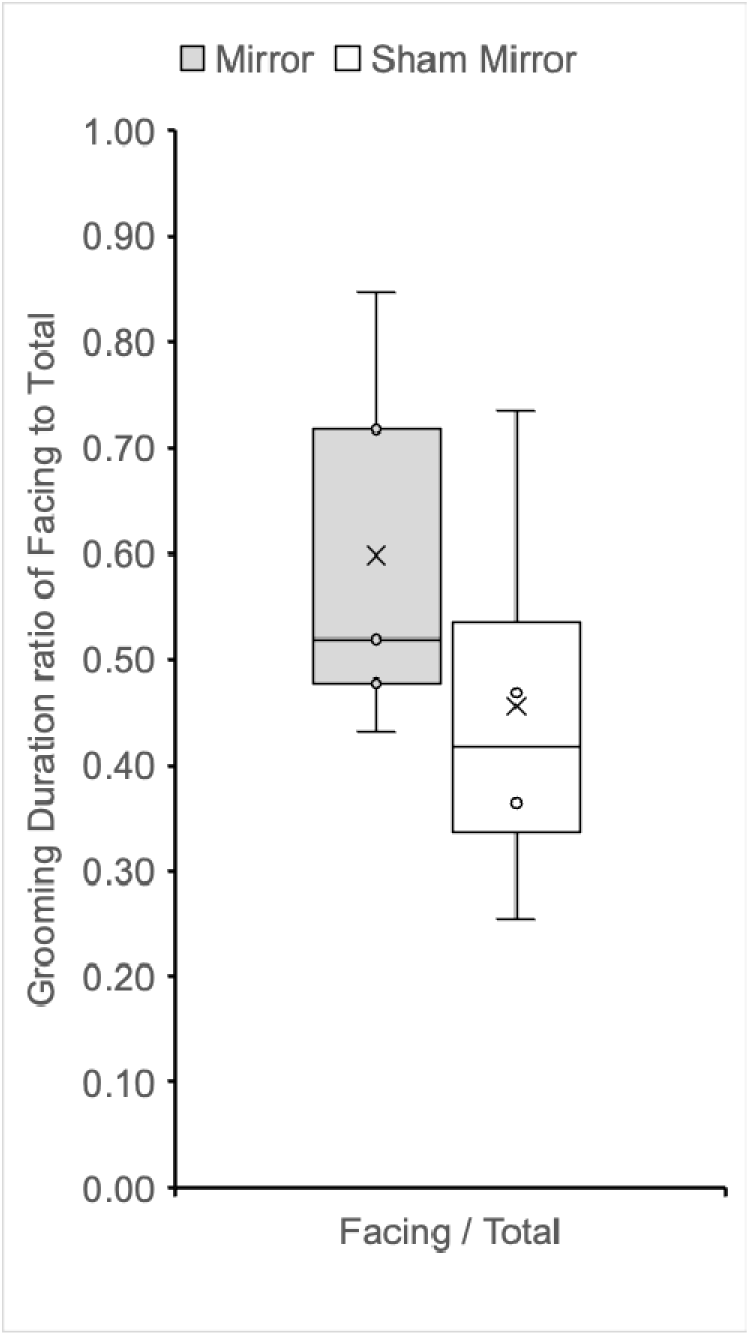
Box plots showing the F/T ratio of grooming duration for the Mirror group and Sham Mirror group in Experiment 2. * * F: Facing the mirror, T: Total.

**Fig 11.**
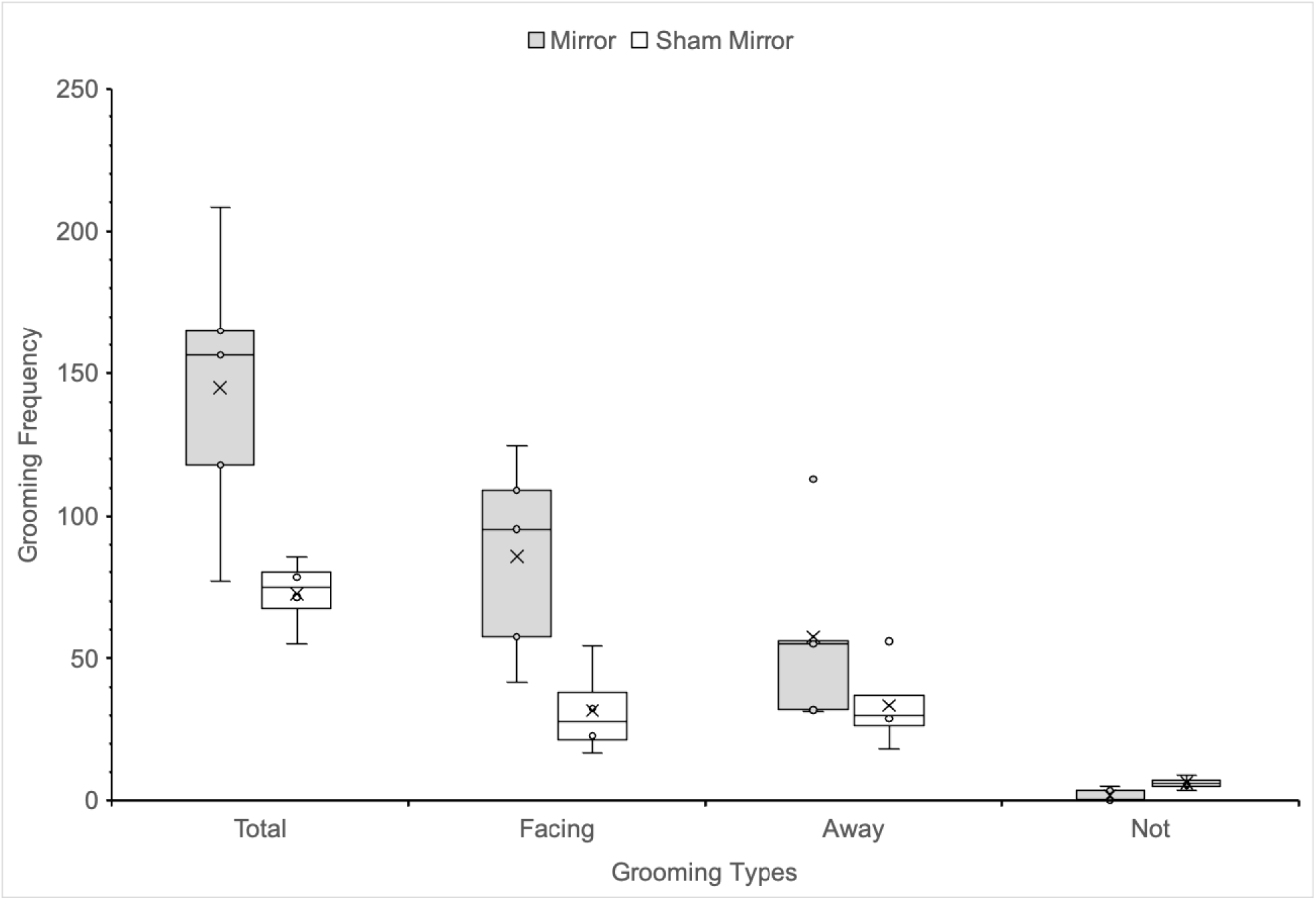
Box plots showing grooming frequency (counts) for T, F, A, and N for the Mirror group and Sham Mirror group in Experiment 2. * * T: Total, F: Facing the mirror, A: Away from the mirror, N: Not facing or away.

**Fig 12.**
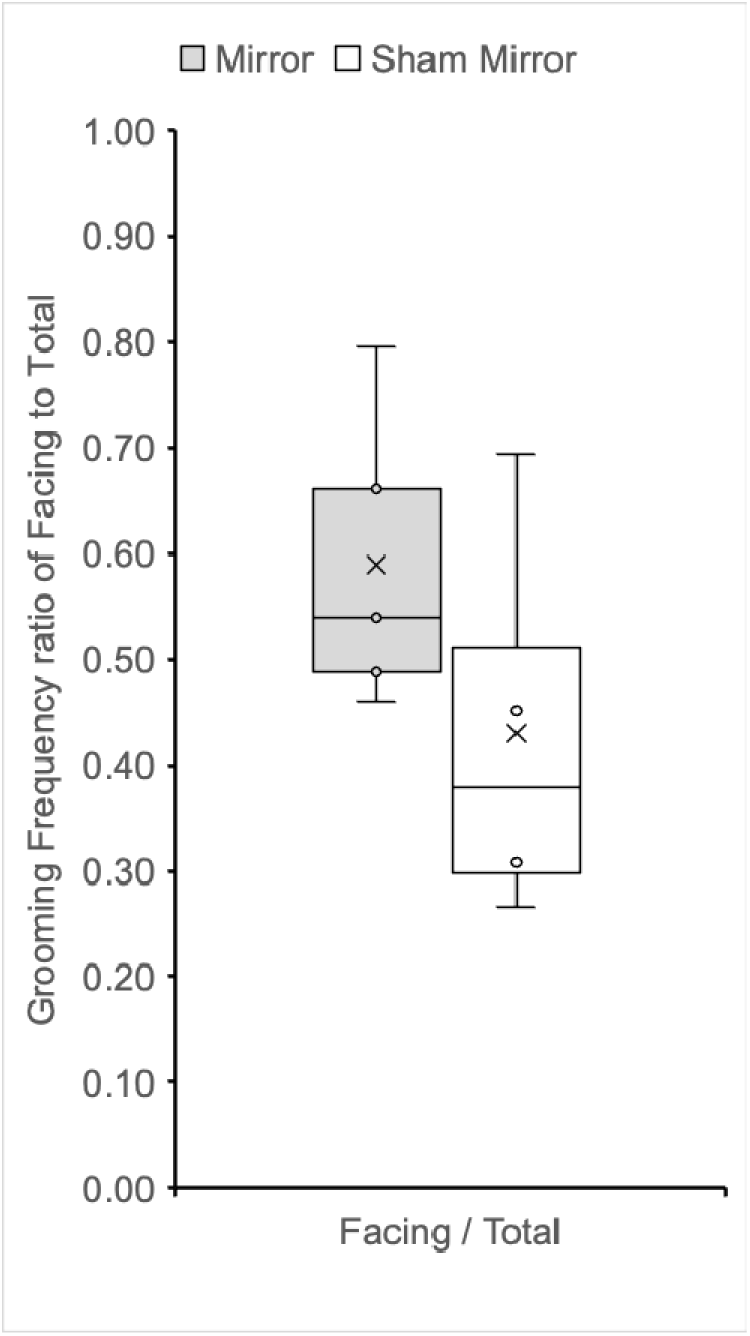
Box plots showing the F/T ratio of grooming frequency for the Mirror group and Sham Mirror group in Experiment 2. * * F: Facing the mirror, T: Total.

### Grooming duration

Within-group (Mirror, n = 5): No significant difference between mean duration Facing (M = 16.951, SD = 8.140) and Away (M = 11.399, SD = 9.017) from the mirror (Paired t(4) = 1.110, p =.329, d = 0.496, 95% CI for mean difference [-8.334, 19.438], mean difference = 5.552 seconds). Within-group (Sham Mirror, n = 4): No significant difference between mean duration Facing (M = 5.741, SD = 2.852) and Away (M = 5.888, SD = 3.613) from the sham mirror (Paired t(3) =-0.050, p =.963, d =-0.025, 95% CI for mean difference [-9.369, 9.077], mean difference =-0.146 seconds).

Between-group (Total): Significantly longer duration in the Mirror group (M=28.549, SD = 12.839) than the Sham Mirror group (M = 12.748, SD = 2.806) (Welch’s t(4.471) = 2.673, p =.050, d = 1.600, 95% CI for mean difference [0.051, 31.553], mean difference = 15.802 seconds). Between-group (Facing): Significantly longer duration Facing the mirror in the Mirror group (M = 16.951, SD = 8.140) than the Sham Mirror group (M = 5.741, SD = 2.852) (Welch’s t(5.160) = 2.867, p =.034, d = 1.743, 95% CI for mean difference [1.253, 21.167], mean difference = 11.210 seconds). Between-group (Away): No significant difference between Mirror (M = 11.399, SD = 9.017) and Sham Mirror (M = 5.888, SD = 3.613) groups (Welch’s t(5.473) = 1.247, p =.263, d = 0.764, 95% CI for mean difference [-5.558, 16.581], mean difference = 5.512 seconds). Between-group (Facing/Total Ratio): No significant difference between Mirror (M =.598, SD = 0.177) and Sham Mirror (M =.456, SD = 0.206) groups (Welch’s t(6.008) = 1.100, p =.314, d = 0.752, 95% CI for mean difference [-0.175, 0.460], mean difference =.143).

### Grooming frequency

Within-group (Mirror, n = 5): A non-significant trend for more frequent grooming Facing (M = 85.600, SD = 34.976) than Away (M = 57.500, SD = 33.227) from the mirror (Paired t(4) = 1.413, p =.231, d = 0.632, 95% CI for mean difference [-27.125, 83.325], mean difference = 28.100). Within-group (Sham Mirror, n = 4): No significant difference between frequency Facing (M = 31.750, SD = 16.454) and Away (M = 33.625, SD = 15.892) from the sham mirror (Paired t(3) =-0.129, p =.905, d =-0.065, 95% CI for mean difference [-47.967, 44.217], mean difference =-1.875). Between-group (Total): Significantly higher frequency in the Mirror group (M = 145.000, SD = 49.793) than the Sham Mirror group (M = 72.625, SD = 13.066) (Welch’s t(4.672) = 3.119, p =.029, d = 1.875, 95% CI for mean difference [11.438, 133.312], mean difference = 72.375).

Between-group (Facing): Significantly higher frequency Facing the mirror in the Mirror group (M = 85.600, SD = 34.976) than the Sham Mirror group (M = 31.750, SD = 16.454) (Welch’s t(5.916) = 3.047, p =.023, d = 1.886, 95% CI for mean difference [10.455, 97.245], mean difference = 53.850). Between-group (Away): No significant difference between Mirror (M = 57.500, SD = 33.223) and Sham Mirror (M = 33.625, SD = 15.892) groups (Welch’s t(5.965) = 1.417, p =.207, d = 0.878, 95% CI for mean difference [-17.412, 65.162], mean difference = 23.875). Between-group (Facing/Total Ratio): No significant difference between Mirror (M =.589, SD = 0.139) and Sham Mirror (M =.430, SD = 0.193) groups (Welch’s t(5.318) = 1.383, p =.222, d = 0.966, 95% CI for mean difference [-0.131, 0.449], mean difference =.159).

### Summary for Experiment 2

In Experiment 2, bees with prior mirror exposure (Mirror group) exhibited significantly higher total grooming duration and frequency, and significantly more grooming bouts facing the mirror, compared to the Sham Mirror group. Specifically, the Mirror group showed a significant increase in grooming bouts facing the mirror (Facing), but no significant difference in grooming bouts away from the mirror (Away). However, there was no significant difference in the Facing/Total ratio between the groups. Importantly, however, this increase in grooming did not result in a statistically significant bias facing the mirror when compared to grooming away from it within the Mirror group, nor did the proportion of mirror-facing grooming differ significantly between groups.

### Individual analysis (binomial test)

Directional bias in grooming frequency at the individual level was assessed by comparing F versus A frequencies for each observer per bee (two-tailed binomial test, α =.05).

### Experiment 1: Mirror group

For the Mirror group (n = 6), only S5 showed a significant Facing > Away bias according to both observers (ps <.05). S1 showed this bias only for observer MM. No significant bias was found for the other 4 bees by either observer.

### Experiment 1: Control group

For the Control group (n =6), while some subjects showed significant biases (S3 (RA), S11 (MM) showed Facing > Away), no individual showed a consistent directional bias according to both observers.

### Experiment 2: Mirror group

For the Mirror group (n = 5), S1 and S6 showed a significant Facing > Away bias according to both observers (ps <.05). No significant bias was found for the other 3 bees by either observer.

### Experiment 2: Sham Mirror group

For the Sham Mirror group (n = 4), S3 showed a significant Facing > Away bias according to both observers (ps <.05). S8 showed a significant Away > Facing bias according to both observers (ps <.05). S7 showed an Away > Facing bias only for observer MM. The remaining bee (S5) showed no significant bias according to either observer.

### Summary of individual results

These individual-level analyses reveal that while group averages showed some significant effects (especially in Exp 2), only a minority of subjects in the mirror groups consistently directed significantly more grooming facing the mirror than away (1/6 or ∼17% in Exp 1; 2/5 or 40% in Exp 2, based on agreement between observers). Furthermore, the presence of significant biases (both facing and Away) in some control/sham mirror subjects complicates simple interpretation based solely on directional preference. These findings highlight pronounced individual variation and suggest factors beyond MSR could influence the results, highlighting the need to look beyond simple group averages when evaluating MSR potential.

## Discussion

This study investigated the potential for mirror self-recognition (MSR) in the western honeybee using the classic mark test paradigm. The core logic relied on comparing head-directed grooming directed at a mark (invisible without a mirror) between subjects with and without mirror access, thereby controlling for the baseline effects of the mark itself. Any significant increase in mark-directed grooming specifically in the mirror condition would suggest recognition of the marked self-image.

### Interpretation of findings

#### Preliminary insights requiring cautious interpretation

Our findings provide preliminary insights into how honeybees respond to mirrors after being marked, but the assessment regarding MSR potential requires considerable caution. The results of Experiment 1 particularly highlight this need for caution. In this experiment, a key indicator of MSR was absent, as the mirror group showed no significant increase in the total amount of grooming compared to the control group. And yet, the proportion of grooming directed towards the mirror (F/T ratio) was significantly higher for both duration and frequency. This shift in proportion, without an increase in the total quantity of grooming, can be explained by a simpler mechanism than MSR. Specifically, it is possible that the bees exhibited an orienting response to the mirror as a novel visual stimulus. This well-documented behavior, in which an organism’s attention is captured by a new object, would explain why they spent more time oriented in its direction. Consequently, the baseline grooming behavior—likely prompted by the physical irritation of the mark—was more likely to be observed coincidentally while the bee was facing the mirror. Furthermore, a mirror inherently presents a highly symmetrical visual environment. Previous research has demonstrated that honeybees can extract bilateral symmetry as a visual feature and exhibit a predisposition to maintain closer proximity to and display prolonged visual inspection of symmetrical patterns compared to asymmetrical ones [51]. Therefore, the elevated F/T ratio observed in our study might not reflect self-directed attention, but rather an innate or learned orienting response driven by the low-level visual extraction of the mirror’s symmetrical properties. This explanation, which favors a simpler, non-cognitive explanation, is consistent with findings in other complex invertebrates. For instance, in a preliminary mark test on the common octopus (*Octopus vulgaris*), subjects were found to frequently explore the mark with their arms even in control trials without a mirror. The study concluded that these mark-directed behaviors were likely driven by proprioceptive stimuli from the mark itself, rather than by visual feedback from the mirror [20].

This alternative hypothesis is strongly supported by the finding that even within the mirror group, there was no clear and significant preference for grooming while facing the mirror over grooming while facing away. Therefore, the elevated F/T ratio cannot be conclusively construed as a cognitive leap of self-recognition and may instead be an artifact of the spatial overlap between attentional orientation and a pre-existing grooming pattern.

In contrast, Experiment 2 revealed a different behavioral pattern after mirror pre-exposure. Here, bees in the mirror group exhibited significantly higher total grooming activity than the sham-mirror controls. While this increase might initially seem more consistent with MSR, other key details suggest a different conclusion, as emphasized in critical reviews of MSR research [13]. Crucially, the significant bias in the F/T ratio observed in Experiment 1 disappeared in Experiment 2. Furthermore, as in Experiment 1, bees in the mirror group still did not groom significantly more when facing the mirror compared to when facing away. This implies that while pre-exposure caused the mirror to become a trigger for increased grooming, this behavior was not visually guided by the reflection. Instead, the mirror image may have been perceived as a social stimulus—that is, a conspecific. From this perspective, the increased grooming is not an act of self-recognition, but rather a sign of the resulting heightened arousal, stress, or a social response to this stimulus [e.g., 2, 9, 11, 60].

Taken together, the results from both experiments do not contradict each other; rather, they form a consistent narrative that argues against MSR in honeybees. The initial reaction to the mirror (Experiment 1) appears to be an attentional artifact, while the subsequent reaction after mirror pre-exposure (Experiment 2) appears to be one of generalized agitation. In neither case did the bees display the key criterion for MSR: the specific, tool-like use of the mirror to guide actions toward an otherwise invisible part of the body [e.g., 1-3]. Ultimately, determining whether an animal perceives its reflection as self, another individual, or simply a novel visual stimulus remains a central challenge in MSR research [10, 11]. Our results reinforce the interpretation that the observed behaviors can be explained by simpler, non-cognitive mechanisms. This supports a foundational debate in comparative psychology, where behaviors often attributed to higher-order cognition, such as a’Theory of Mind,’ have been compellingly reinterpreted as the products of associative learning [66], and is a recurring theme in insect cognition where seemingly abstract abilities have been alternatively explained by lower-level processes. For instance, sequential inspection strategies can account for numerosity [67] and spatial tasks [42], while graded perceptual similarity explains matching and generalization [30, 51, 52]. Furthermore, simple neural network models successfully replicate these seemingly complex concept learnings [54, 55]. A prime example of this debate is the finding that honeybees can “opt-out” of difficult discrimination tasks, a behavior seen as evidence for metacognition or “uncertainty monitoring” in primates. Even in this case, however, the authors rigorously discuss the possibility that this seemingly complex behavior could be explained by simpler associative mechanisms rather than a true awareness of uncertainty [45].

### Lack of clear within-group difference (facing vs. away)

A key observation requiring careful consideration is the general lack of a statistically significant preference for grooming while facing the mirror versus grooming while facing away from it within the mirror group in both experiments, particularly in Experiment 2 where overall grooming increased. Classic MSR, as demonstrated in chimpanzees [3], involves the animal specifically using the mirror to direct actions towards the mark. The absence of a clear bias towards mirror-facing grooming for the mark itself in honeybees weakens the MSR interpretation. This is consistent with findings suggesting that some seemingly complex cognitive tasks in bees, such as the learning of’above/below’ spatial relationships [37], may be solved by employing stereotyped flight movements and sequential inspection that transform the problem into a simpler discrimination task [42]. Similarly, numerosity estimation has been explained by a simple neural network coupled with a sequential inspection strategy [67]. These precedents suggest the observed grooming could be a byproduct of a simpler behavioral mechanism rather than genuine self-recognition.

### Substantial individual variation

The pronounced individual variation revealed by both group-level statistics and binomial tests further complicates interpretation. In both experiments, only a minority of subjects in the mirror groups consistently directed significantly more grooming towards the mirror. Moreover, the occurrence of directional biases in some Control/Sham mirror subjects suggests that factors unrelated to MSR could influence the results. This heterogeneity highlights the need to look beyond simple group averages when assessing the possibility of MSR and reinforces the importance of analyzing individual response patterns [4, 13].

### Alternative interpretations

#### Conspecific or novel stimulus recognition

A plausible alternative interpretation, as often discussed in MSR literature [11], is that bees might have responded to the reflection as if it were a novel bee, a familiar bee with an unusual mark, or simply an unfamiliar visual stimulus. This type of social responding to a mirror image has been clearly demonstrated in other invertebrates. For example, female cuttlefish (*Sepia officinalis*) display a specific body pattern called’Splotch’ almost exclusively towards other females and their own mirror image, suggesting they perceive their reflection as a same-sex conspecific [68]. This principle of seeking simpler, more parsimonious explanations is a foundational tenet of comparative psychology, famously expressed as Morgan’s Canon [69], and is echoed in the rigorous analytical framework required for comparing diverse species, which cautions against inferring complex cognitive processes when simpler mechanisms cannot be ruled out [65, 66]. Indeed, the challenge of disentangling abstract conceptual understanding from reliance on lower-level mechanisms, such as graded perceptual similarity, has been a central debate in the field ever since the pioneering work on relational matching in primates [70]. A prior investigation into same-different learning argued that bees’ performance in visual matching tasks could instead be explained by reliance on lower-level mechanisms such as graded perceptual similarity or other simple visual cues [30], an argument similarly applied to their ability to extract visual symmetry [51]. Applying this same critical lens to the current study, it is imperative to consider that the observed mirror-induced grooming may not reflect the cognitive leap of self-recognition, but rather a manifestation of these more fundamental behavioral or perceptual responses. This misinterpretation could trigger various behaviors such as increased antennation and touching, potentially alongside social, defensive, or exploratory responses. Notably, a similar study on paper wasps [17] found increased exploratory behaviors (antennation and touching) but a decrease in self-directed grooming in response to mirrors, suggesting these behaviors may serve different functions across insect species and not necessarily all point towards MSR. The honeybee’s grooming could be a displacement activity triggered by the ambiguous stimulus of its reflection [16], rather than a mark-directed action guided by self-recognition. The phenomenon of social responding to mirrors, even after extensive habituation, is well-documented in primates such as rhesus monkeys, where a simple change in the mirror’s location can reinstate social responses [71], and notably, self-recognition is absent in social pairings like mother-infant or infant-infant groups [72].

The current study design, lacking a crucial control condition where bees are presented with a live, marked conspecific, makes it difficult to definitively distinguish this social/novelty-response hypothesis from the MSR hypothesis.

### Methodological prerequisites and the mark test

Beyond cognitive capacity, recent critiques emphasize that an animal’s performance on the mark test is contingent on fulfilling several methodological prerequisites [73]. First, the animal must understand the properties of a mirror. Our pre-exposure in Experiment 2 was designed to facilitate this, and the resulting increase in grooming suggests the mirror had an effect, but it is uncertain if this constitutes true mirror understanding.

Second, the mark itself must be salient enough to motivate investigation. While the white paint was visually conspicuous to humans, its motivational salience for a honeybee, which relies heavily on tactile and olfactory cues for grooming, is questionable. Third, the animal must possess a repertoire of self-exploratory behavior. While bees groom, it is unknown if this extends to visually-guided exploration of a specific, marked body part. The failure to meet one or more of these criteria, as outlined by Kakrada & Colombo (2022) [73], could explain our inconclusive results, independent of whether honeybees possess a self-concept.

### The perceptual-motor skill hypothesis

An even more parsimonious explanation is that MSR-like behavior may not reflect an abstract self-concept at all, but rather a learned perceptual-motor skill [74]. Landmark studies demonstrated that pigeons, through explicit operant conditioning that links seeing a mark in the mirror with pecking their own body, can be trained to pass the mark test [75, 76]. From this perspective, the increased grooming we observed in Experiment 2 could be a low-level associative response to the novel visuotactile contingency created by the mark and the mirror, rather than an act of self-recognition.

This critical stance is warranted even when bees demonstrate robust cognitive abilities under rigorous conditions. For instance, even when honeybees showed evidence of concept learning using more rigorous methods, such as a trial-unique procedure designed to minimize simpler associative explanations, the potential for alternative, lower-level mechanisms still required careful consideration [30]. This aligns with our conclusion that seemingly complex behaviors can often be explained by simpler mechanisms, a principle rigorously debated across taxa. For instance, while primates and avians have historically been the focus of complex cognition [66, 75, 76], recent reviews emphasize that seemingly abstract conceptualizations can also be achieved by the miniature brains of insects [77], often through simpler, lower-level associative mechanisms [67, 78].

### Mark/anesthesia effects

While the comparison design aimed to control for the effects of the marking and anesthesia procedures, their potential influence as confounding variables cannot be entirely dismissed. For instance, the physical irritation or olfactory cues from the marking fluid itself may have increased baseline grooming across all conditions. Future research should incorporate dedicated no-mark control groups to fully isolate these potential effects [1, 2, 10]. Consequently, the observed grooming behavior—even when performed while facing the mirror—may not be a visually-guided attempt to touch the mark, but rather a generalized response to the physical irritation or olfactory cues from the marking fluid itself. The antennae are vital sensory organs, and any irritation nearby could easily trigger cleaning behaviors directed at them, independent of any visual information from the mirror.

### Non-specific mirror effects

The vertical mirror might induce spatial confusion or general arousal unrelated to self-recognition [cf. 62, 63]. Honeybees are known to be highly sensitive to visual environmental changes [46].

### Methodological limitations

Several methodological limitations must be considered when interpreting our findings.

Primarily, the small sample sizes (n=4−6 per group in Experiment 1 after exclusions, and n=4−5 per group in Experiment 2) result in low statistical power, which significantly affects the ability to draw firm conclusions. Additionally, the observers were not blind to the experimental conditions, which introduces a potential for unconscious observer bias.

Perhaps the most significant methodological limitation, central to the interpretation of all our findings, was the ambiguity in our primary dependent variable: the observers’ inability to reliably distinguish between grooming actions specifically targeting the mark and more generalized head-grooming (e.g., of the antennae). This means we cannot definitively conclude that the observed grooming, even when performed while facing the mirror, was a contingent response to *seeing* the mark. Instead, it is highly plausible that the underlying motivation for grooming stems from non-visual stimuli (e.g., tactile irritation or olfactory cues) from the marking fluid itself. In this view, the mirror does not serve as a tool for guided self-inspection, but rather as an arousing visual stimulus that triggers an increase in this pre-existing, generalized grooming behavior. Because our methodology could not falsify this simpler, non-cognitive hypothesis, the data do not allow us to confirm that the bees were using the mirror as a tool for targeted self-inspection. Given all these limitations, the statistical findings should be interpreted with extreme caution, and we stress the necessity of future replications with larger sample sizes and refined methodologies.

However, as has been argued for cephalopods, a failure to pass the classic mark test does not necessarily confirm the absence of self-awareness [18]. The interpretation of mirror-induced behaviors must consider the specific sensory world of the animal in question. Honeybees [23, 24, 26], much like cuttlefish, are sensitive to polarized light. Research on cuttlefish has shown that they react differently to their mirror image depending on whether the reflection’s polarization patterns are intact, suggesting they are perceiving subtle cues in their reflection that go beyond simple shape and color [79]. This highlights the possibility that insects may also perceive their reflections in ways not immediately obvious to human observers, adding another layer of complexity to the interpretation of the mark test.

### Comparison with other invertebrate studies

Our preliminary findings, particularly the increased grooming in Experiment 2, contrast somewhat with studies on paper wasps [17] that found no clear MSR evidence, although they reported increased exploratory behaviors like antennation and touching the mirror. The wasp study employed crucial control conditions absent here (e.g., sham-marked wasps with a mirror). The findings highlight the sensitivity of wasps to mirror presence and marking, suggesting they perceive these alterations but respond differently than classic MSR would predict (e.g., with more exploration rather than self-directed grooming). Their results emphasize the difficulty in demonstrating conclusive MSR in insects using this paradigm. Our results add to the sparse literature on potential MSR in invertebrates like ants [16] and crabs [21], all of which present interpretational challenges. While evidence for MSR in cephalopods is also contested, recent work using a different paradigm—the rubber arm illusion—suggests that octopuses possess a sophisticated sense of body ownership by integrating visual, tactile, and proprioceptive information [80]. This finding indicates that complex forms of self-perception may exist in invertebrates, which the classic mark test might not be suited to detect [80]. Further supporting this notion, recent studies on bumblebees have shown that they perceive the affordance of narrow gaps in relation to their own body size, turning sideways to avoid collisions in a manner strikingly similar to humans [81]. This capacity to gauge the relationship between the environment and one’s own body dimensions suggests a form of bodily self-awareness that operates independently of mirror recognition [82].

### Broader implications and future directions

If honeybees possess even a rudimentary form of MSR—a possibility that remains to be rigorously tested—it would significantly challenge assumptions about the cognitive and neural requirements for this ability, contributing to the broader debate on whether larger brains are necessarily “better” [8], and aligns with the emerging perspective that the neural substrates for subjective experience, a foundational aspect of self-awareness, may not be exclusive to vertebrates. Indeed, it has been compellingly argued that the insect brain possesses the necessary functional organization to support a capacity for basic consciousness [83]. It would support a “gradualist perspective” on self-awareness, suggesting that this capacity is not an all-or-nothing phenomenon but rather exists on a continuum across the animal kingdom [14, 17, 21], ranging from fundamental bodily self-perception [21, 74, 80, 83] to more abstract self-concepts [9, 13, 74, 83]. This finding would reinforce the idea that complex behaviors can arise from relatively small nervous systems [8, 28, 30, 31, 78], a notion corroborated by computational modeling studies demonstrating that abstract concept learning can be achieved with simple neural networks inspired by the insect brain [54] and other computational frameworks [55], and even that numerosity estimation can be enabled by an artificial network of just four neurons [67]. The prerequisite ability for honeybees to generalize visual patterns to their mirror images [52] provides a fascinating cognitive foundation, suggesting that while bees can process mirror-image information, this does not automatically translate to recognizing their own image.

However, given the significant limitations and strong critiques leveled against non-primate MSR claims [13], firm conclusions are premature. The history of MSR research includes numerous claims that did not stand up to rigorous scrutiny, such as initial failures to find MSR in great apes like gorillas, or early studies on monkeys that lacked adequate controls [1, 2, 10]. Indeed, a central critique within the primate literature itself is that even when chimpanzees pass the test, their behavior can be interpreted as a sophisticated understanding of the mirror’s properties for guiding the body, rather than as evidence for a human-like, introspective self-concept [11]. This history provides a critical lesson for invertebrate research. As Heyes argued in her review of the primate literature, the burden of proof for complex cognitive claims lies in experimentally ruling out simpler, non-mentalistic alternative explanations [66]. Therefore, future research on honeybees should not only aim for replication but must incorporate the rigorous control conditions necessary to falsify these leaner hypotheses. Specifically, this requires studies designed with: (1) increased statistical power through larger sample sizes, (2) the implementation of essential control groups (e.g., no-mark, conspecific controls), (3) refined behavioral analysis, and (4) further investigation of individual differences.

## Conclusion

This exploratory study employed a mark test to investigate MSR potential in honeybees. Experiment 2 (with mirror pre-exposure) showed increased overall grooming, more likely attributable to generalized arousal than specific self-recognition. Our findings must be considered preliminary due to the lack of a significant preference for mirror-facing grooming within the mirror group, substantial individual variability, plausible alternative explanations, and critical methodological limitations, including low statistical power. The results do not provide conclusive evidence for MSR capability in honeybees at this stage. However, a failure to pass the classic mark test does not necessarily confirm the absence of self-awareness, particularly when considering the specific sensory world of the animal in question [10, 14, 18, 20, 21, 57, 74, 79].

Ultimately, our preliminary findings underscore a critical principle in comparative psychology, echoing the tenet of Morgan’s Canon [69] and the modern intellectual caution that demands the rigorous exclusion of simpler explanations before accepting complex cognitive claims [45, 64, 66]. Therefore, while our results do not provide conclusive evidence for MSR, their primary value lies in highlighting specific and critical avenues for future investigation. Building upon the framework established here, future research should not only prioritize increasing statistical power but also incorporate refined control conditions to directly address the alternative hypotheses discussed, such as testing for mark salience, distinguishing MSR from social responses, and designing paradigms that can disentangle associative learning from a potential self-concept.

Furthermore, exploring alternative paradigms that may reveal different facets of self-perception, such as the bodily self-awareness demonstrated in other invertebrates [18, 20, 21, 80–82], will be essential. By systematically exploring these issues, future work can clarify the cognitive limits of the honeybee mind and its place in the broader debate on the convergent evolution of cognition [25] and self-awareness.

## Supporting information

S1_Experiment1_RawData

S2_Experiment2_RawData

S3_Experiment2_MirrorGroup(S6)_GroomingVideo_VideoS1

## Acknowledgments

The authors acknowledge the use of Gemini 3.1 pro for assistance with language editing and improving the clarity of the manuscript. The authors reviewed and take full responsibility for the final content.

## Funding

The authors received no specific funding for this work. All research materials and costs were supported by institutional funds from Mita International School of Science. The funders had no role in study design, data collection and analysis, decision to publish, or preparation of the manuscript.

## Author contributions

Conceptualization: Gentaro Shishimi

Data curation: Gentaro Shishimi, Rena Akaishi, Mana Matsukami

Formal analysis: Gentaro Shishimi

Investigation: Rena Akaishi, Mana Matsukami

Methodology: Gentaro Shishimi

Project administration: Gentaro Shishimi

Supervision: Gentaro Shishimi

Visualization: Gentaro Shishimi, Rena Akaishi, Mana Matsukami

Writing – original draft: Gentaro Shishimi, Rena Akaishi, Mana Matsukami

Writing – review & editing: Gentaro Shishimi, Rena Akaishi, Mana Matsukami

## Data availability statement

The raw data for Experiment 1 and Experiment 2 are provided as Supporting Information files S1 and S2, respectively. A video illustrating head-directed grooming behavior of S6 in the mirror group of Experiment 2 is available as Supporting Information file S3 (Video S1).

## Notes

### Competing Interest Statement

The authors have declared no competing interest.

### Summary of Updates

This revised version (v2) includes the following updates and improvements: Expanded Discussion: We further explored alternative interpretations of the observed mirror-induced behaviors. Additional literature (e.g., Giurfa et al., 1996; Perry & Barron, 2013) was incorporated to discuss how symmetry perception and metacognition-like behaviors in honeybees can be explained by lower-level associative mechanisms, thereby strengthening the perceptual-motor skill hypothesis. Citation Reorganization: The in-text citations and the reference list were comprehensively reorganized and unpacked to improve clarity, ensuring precise attribution of specific cognitive processes to their respective studies. Figure and Formatting Updates: The box plots (Figures 5-12) were reordered to strictly match the sequence discussed in the text (Total, Facing, Away, Not facing/away). Minor typographical errors were corrected, graph axis labels were capitalized (e.g., 'types' to 'Types'), paragraph breaks were adjusted for better flow, and methodological descriptions regarding the test box dimensions were refined. Acknowledgments: Updated the AI tool version referenced for language editing assistance.

